# Spatially varying cis-regulatory divergence in *Drosophila* embryos elucidates cis-regulatory logic

**DOI:** 10.1101/175059

**Authors:** Peter A. Combs, Hunter B. Fraser

## Abstract

Spatial patterning of gene expression is a key process in development—responsible for the incredible diversity of animal body plans—yet how it evolves is still poorly understood. Both cis- and trans-acting changes could accumulate and participate in complex interactions, so to isolate the cis-regulatory component of patterning evolution, we measured allele-specific spatial gene expression patterns in *D. melanogaster* × *D. simulans* hybrid embryos. RNA-seq of cryosectioned slices revealed 55 genes with strong spatially varying allele-specific expression, and several hundred more with weaker but significant spatial divergence. For example, we found that *hunchback (hb)*, a major regulator of developmental patterning, had reduced expression specifically in the anterior tip of *D. simulans* embryos. Mathematical modeling of *hb* cis-regulation suggested that a mutation in a Bicoid binding site was responsible, which we verified using CRISPR-Cas9 genome editing. In sum, even comparing morphologically near-identical species we identified a substantial amount of spatial variation in gene expression, suggesting that development is robust to many such changes, but also that natural selection may have ample raw material for evolving new body plans via cis-regulatory divergence.

## Introduction

Although most cells in any metazoan share the same genome, they nevertheless diversify into an impressive variety of precisely localized cell types during development. This complex spatial patterning is due to the precise expression of genes at different locations and times during development. Where and when each gene is expressed is largely dictated by the activities of cis-regulatory modules (CRMs, also sometimes called enhancers) through the binding of transcription factors to their recognition sequences (***Banerji et al., 1981***; ***Ptashne, 1986***; ***Driever et al., 1989***). Despite the importance of these patterning CRMs for proper organismal development, they are able to tolerate some modest variation in sequence and level of activity (***Ludwig and Kreitman, 1995***; ***Lusk and Eisen, 2010***; ***Villar et al., 2015***; ***Berthelot et al., 2017***). Indeed, this variation is one of the substrates upon which selection can act. However, even in the handful of cases where we understand the regulatory logic, efforts to predict the result of inter-specific differences in CRMs still have limited precision (***Small et al., 1991***; ***Samee and Sinha, 2014***; ***Sayal et al., 2016***).

A complicating factor in comparing gene expression between species is that both cis- and trans-acting regulation can change (***Coolon et al., 2014***). One solution is to focus on cis-regulatory changes by measuring allele-specific expression (ASE) in F1 hybrids. In a hybrid each diploid nucleus has one copy of each parent’s genome which is exposed to the same trans-environment, so any differences in zygotic usage of the two copies is due either to cis-regulatory divergence or to stochastic bursting (which should be averaged out over many cells). The early *Drosophila* embryo provides a unique opportunity to probe the interaction of trans-regulatory environments with cis-regulatory sequence: by slicing the embryo along the anterior-posterior axis, we are able to measure ASE in nuclei with similar complements of transcription factors (TFs). By combining knowledge of both the regulatory sequence changes between the species and the transcription factors expressed in each slice, it should be possible to more quickly identify which TF binding site underlies the expression difference.

In this study, we used spatially-resolved transcriptome profiling to search for genes where cis-regulatory differences drive allele-specific expression patterns in hybrid *D. melanogaster*◊*D. simulans* embryos (specifically the reference strains DGRP line 340 for *D. melanogaster* and *w*^501^ for *D. simulans*; we will refer specifically to the two reference strains, and not the two species as a whole unless otherwise noted). We found dozens of genes with clear, consistent differences in allele-specific expression across the embryo. We chose one of these genes, *hunchback (hb)*, as a model to understand which of 17 polymorphisms in its regulatory regions was likely to drive the expression difference. Mathematical modeling of *hunchback* cis-regulation suggested that a Bicoid binding site change was responsible for the expression difference, which we confirmed through CRISPR-Cas9 mediated editing of the endogenous *D. melanogaster* locus.

## Results

### A genome-wide atlas of spatial gene expression in D. melanogaster × D. simulans hybrids

We selected five mid-stage 5 hybrid embryos, with membrane invagination between 50 and 65%. We then sliced the embryos to a resolution of 14µ, yielding between 24 and 27 slices per embryo. We chose embryos from reciprocal crosses (i.e. with either a *D. melanogaster* mother or a *D. simulans* mother), and had at least one embryo of each sex from each direction of the cross. Although hybrid female embryos with a *D. simulans* mother are embryonic lethal at approximately this stage due to a heterochromatin segregation defect (***Ferree and Barbash, 2009***), they were morphologically normal and so we included one female embryo from this cross. We also sliced one embryo from each of the parental strains. Following slicing, we amplified and sequenced poly-adenylated mRNA using SMART-seq2 with minor modifications (***Combs, 2015***; ***Picelli et al., 2014***, ***2013***).

We first searched for cases of hybrid mis-expression—genes where the absolute expression pattern is consistently different in the hybrid, compared to the parents alone. Using earth-mover distance (EMD) to measure differences in expression patterns (***Figure 1—Figure supplement 2A***; ***Rubner et al.*** (***1998***)), for each zygotically expressed gene we compared the expression pattern from each of the hybrid embryos to the pattern expected by taking the average of the *D. melanogaster* and *D. simulans* embryos. After Benjamini-Hochberg FDR correction, no gene was significantly more different from the average of the parental embryos than each of the parental embryos were from each other (smallest q-value =.37, see Methods). We also compared expression patterns between hybrid embryos with a *D. melanogaster* mother to those with a *D. simulans* mother, and found that most differences seemed to be due to differing patterns of maternal deposition or noisy expression (***Figure 1–Figure supplement 3***). Thus, we conclude that there do not seem to be any expression patterns that are not explained by differences in the parents or that are unique to the hybrid context.

**Figure 1.**
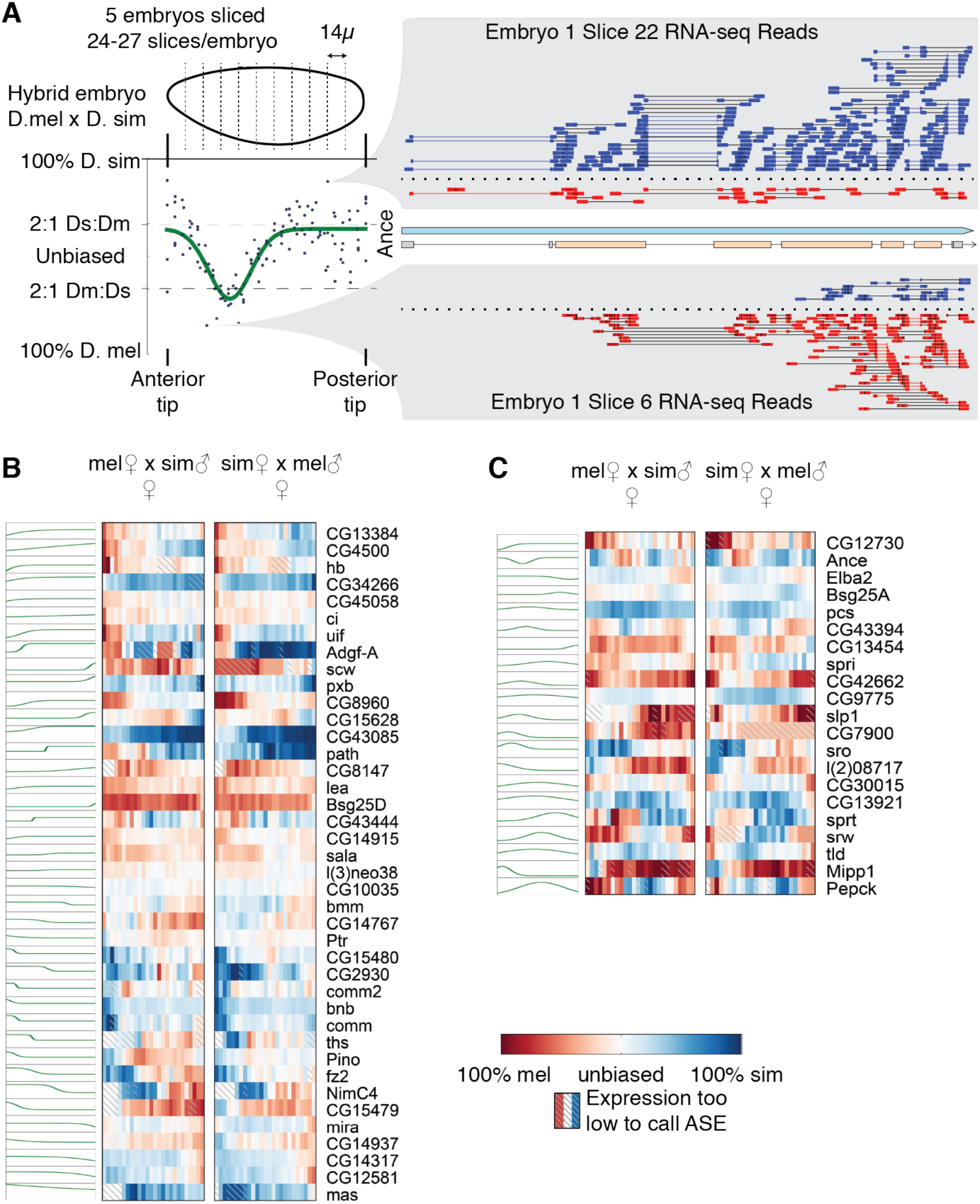
RNA-seq of hybrid *Drosophila* embryos reveals extensive spatially patterned allele-specific expression. A) Each embryo was cryosliced along the anterior-posterior axis in 14µ sections, followed by RNA-seq in each slice. Allele-specific expression (ASE) was called for each gene in each slice by assigning unambiguous reads to the parent of origin; shown here are the reads for the gene *Ance*, with blue indicating *D. simulans* reads and red indicating *D. melanogaster* reads. For each gene, we fit either a step-like or peak-like (shown) function. B-C) Genes with a step-like pattern (B, best fit by a logistic function) or peak-like pattern (C, best fit by a Gaussian function). For each gene, anterior is left and posterior is right. The green line indicates the best fit pattern, with higher indicating *D. simulans* biased expression, and lower indicating *D. melanogaster* biased expression. The heatmaps are from two of the five embryos.

### Overall Allele-specific Expression

In order to measure cis-regulatory differences in expression, we calculated allele-specific expression (ASE) scores for each gene in each slice (***Figure 1A***). The ASE score is the ratio of the difference between the number of *D. simulans* and *D. melanogaster* reads and the sum of the reads,

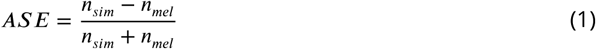

and ranges between -1 (100% *D. melanogaster* expression) and 1 (100% *D. simulans* expression).

Consistent with previous observations (***Wittkopp et al., 2006***; ***Coolon et al., 2012***), we did not find any convincing evidence of imprinting (i.e. zygotic transcription of the maternal or paternal copy of a gene). Although we identified 2,778 genes with a strong maternal expression pattern, defined here as 65% of the slices in all embryos having at least 66.7% of transcripts coming from the mother’s species, these are consistent with the transcripts having been deposited in the egg. Furthermore no genes expressing primarily maternal transcripts had distinct non-uniform expression, consistent with maternal deposition. We also searched for paternally expressed alleles, which would represent strong evidence of imprinting. Because the two-thirds cutoff was quite conservative, we performed separate t-tests on the ASE values in hybrid embryos with *D. melanogaster* mothers and hybrid embryos with *D. simulans* mothers, and took the larger one-sided p-value (reflecting the significance of paternal bias) for each gene. No genes had even a nominal p-value less than 0.1 (i.e. without correcting for multiple testing), suggesting that there are no paternally-biased genes at this stage of development.

Our list of maternally deposited genes is highly concordant with previous measurements of maternal expression. Of the genes classified as maternally expressed in the early expression time-course in ***Lott et al.*** (***2011***), we measured allele-specific expression for 2,653, and found that we clearly agreed on 1,670 (in 552 of the remaining genes, we found the expression to be maternally biased in one of the directions of the cross, but we also detect non-trivial zygotic expression in the other direction). There were also 1,771 maternally provided genes that had low expression (less than 10 FPKM in 65% or more of the slices) in our data, which is consistent with many maternally provided genes being heavily degraded by this point in development. Furthermore, of the 8 genes that ***Lott et al.*** (***2011***) classified as zygotically expressed, we classified as maternally expressed, and which had published *in situ* hybridization data, ***Tomancak et al.*** (***2002***) detected maternally deposited RNA for 5/8, suggesting that they may be dependent on the precise strain or conditions (***Figure 1—Figure supplement 4***). The 564 genes we classified as not biased that ***Lott et al.*** (***2011***) classified as maternal are generally weakly biased as maternal, but not enough to clear our thresholds (***Figure 1—Figure supplement 5***).

We then looked for genes that are consistently biased towards one species, regardless of parent. We found 572 genes (at a 10% FDR) where the overall expression was more biased than expected by chance (see Methods). However, many of these showed only a weak bias (some cases have as few as 2% more reads from one species than from the other), so we further identified a subset of these with at least 2-fold more reads from one species than the other in 65% of slices; we called this subset strongly biased (see Methods). We found 42 genes with strongly *D. melanogaster*-biased expression, and 38 genes with strongly *D. simulans*-biased expression (***Figure 1—Figure supplement 7***). Given that the gene models we are using are taken entirely from *D. melanogaster*, we may be underestimating the true quantity of *D. simulans* biased genes (this caveat does not apply to spatially varying ASE, since inaccurate gene models would not lead to spatial variation across the embryo). Intriguingly, a few of these genes are expressed at comparable levels and with similar spatial patterns in the *D. melanogaster* and *D. simulans* parental embryos, indicating they may be affected by compensatory cis- and trans-acting changes. These species-biased genes are spread throughout the genome, suggesting that this effect is not a consequence of a single cis-regulatory change or inactivation of an entire chromosome.

### Spatially varying allele-specific expression highlights genes with cis-regulatory changes

The greatest power of this dataset lies in its ability to identify genes with spatially varying ASE (svASE)—that is, expression in one part of the embryo that is differently biased than another part of the embryo. In order to identify these genes, we fit two different simple patterns to the ASE as a function of embryo position (***Figure 1A***). We identified 40 genes where a sigmoid function explained at least 45% of the variance in ASE (***Figure 1B***), and 21 where a Gaussian function explained at least that much of the variance (***Figure 1C***; if both explained over half the variance for a gene, we only count the one that better explains the variance). In order to estimate a false discovery rate, we shuffed the *x*-coordinates of the ASE values, and refit the functions. Of 1000 shuffes, only 6 (sigmoid) and 0 (peak) genes cleared the threshold for svASE, which implies false discovery rates of ˘0.020396% (sigmoid) and *<*0.001925% (peak). At a more relaxed 10% FDR cutoff, we found 320 genes where fitting explains at least 12% of the variance in ASE.

We observed very few spatially varying splicing differences in our data (***Figure 1—Figure supplement 8***). In one case, our data suggest that the shorter *A* isoform of the *kni* gene is preferentially expressed in the posterior expression domain; to our knowledge, spatially varying splicing has not been previously observed for *kni*, though the two expression domains are known to be driven by different trans-regulatory factors (***Rothe et al., 1994***). Most examples of spatially varying splice-junction usage qualitatively matched the svASE for the same gene, though it was noisier due to the smaller number of reads supporting splice junction usage compared to expression. An exception to this involved the maternal-zygotic gene *HnRNP-K*, where the shortest isoform was zygotically expressed, consistent with our previous observations that zygotic transcripts are often short in this stage of *Drosophila* development (***Artieri and Fraser, 2014***). The use of alternative first exons in both of these cases suggests that cis-regulation may contribute to the preponderance of short transcripts during early development, in addition to temporal constraints on the transcription of long genes.

Searching for Gene Ontology (GO) function terms enriched for genes with svASE (***Eden et al., 2007***, ***2009***), we found enrichments for genes involved in embryonic morphogenesis (GO:0048598, q-value 2.3 ù 10^−6^), including transcription factors (GO:0003700, q-value 9.8 ù 10^−7^) and transmembrane receptors (GO:0099600, q-value 2.2 ù 10^−2^). These included key components in important signaling pathways, such as *fz2* (a Wnt receptor) and *sog* (a repressor of the TGF-βsignaling pathway). *Myc*,a cell cycle regulator that is a target of both of these pathways, also had significant svASE. However, when we used all non-uniformly expressed genes from ***Combs and Eisen*** (***2013***) as a background set, we did not find any enriched GO terms, suggesting that the enrichments are driven by functions shared by spatially patterned genes overall, rather than among svASE genes specifically.

### A single SNP is the source of svASE in the gap gene *hunchback*

We noticed that *hunchback*, an important transcriptional regulator (***Small et al., 1991***; ***Wimmer et al., 2000***; ***Jaeger, 2011***), had strong svASE (step-like fit *r*^2^ = 0.57; ***Figure 1B***). Since the regulation of *hb* is relatively well-characterized, this provided the opportunity to study the sequence-level causes of the svASE that we observed.

The *hb* svASE was driven by the anterior tip, which had a strong bias towards the *D. melanogaster* allele, suggesting an expansion of the anterior domain relative to *D. simulans* (***Figure 2A***). Compared to ASE elsewhere in the embryo, ASE in the anterior tip was both stronger (Ì 10-fold more *D. melanogaster* transcripts than *D. simulans*), and also less affected by the species of the mother (excluding the first six anterior slices, there are 5-15% more reads from the maternal species than the paternal). When we performed *in situ* hybridization for *hb* RNA, we found overall similar patterns of localization, except in the anterior tip, where we observed *hb* expression in *D. melanogaster*, but not in *D. simulans* (Fig. 2B and C). Although the parental embryos are not precisely the same size, the *in situs* are consistent with the svASE, suggesting that the divergence is not due to embryo size or trans-regulatory changes.

**Figure 2.**
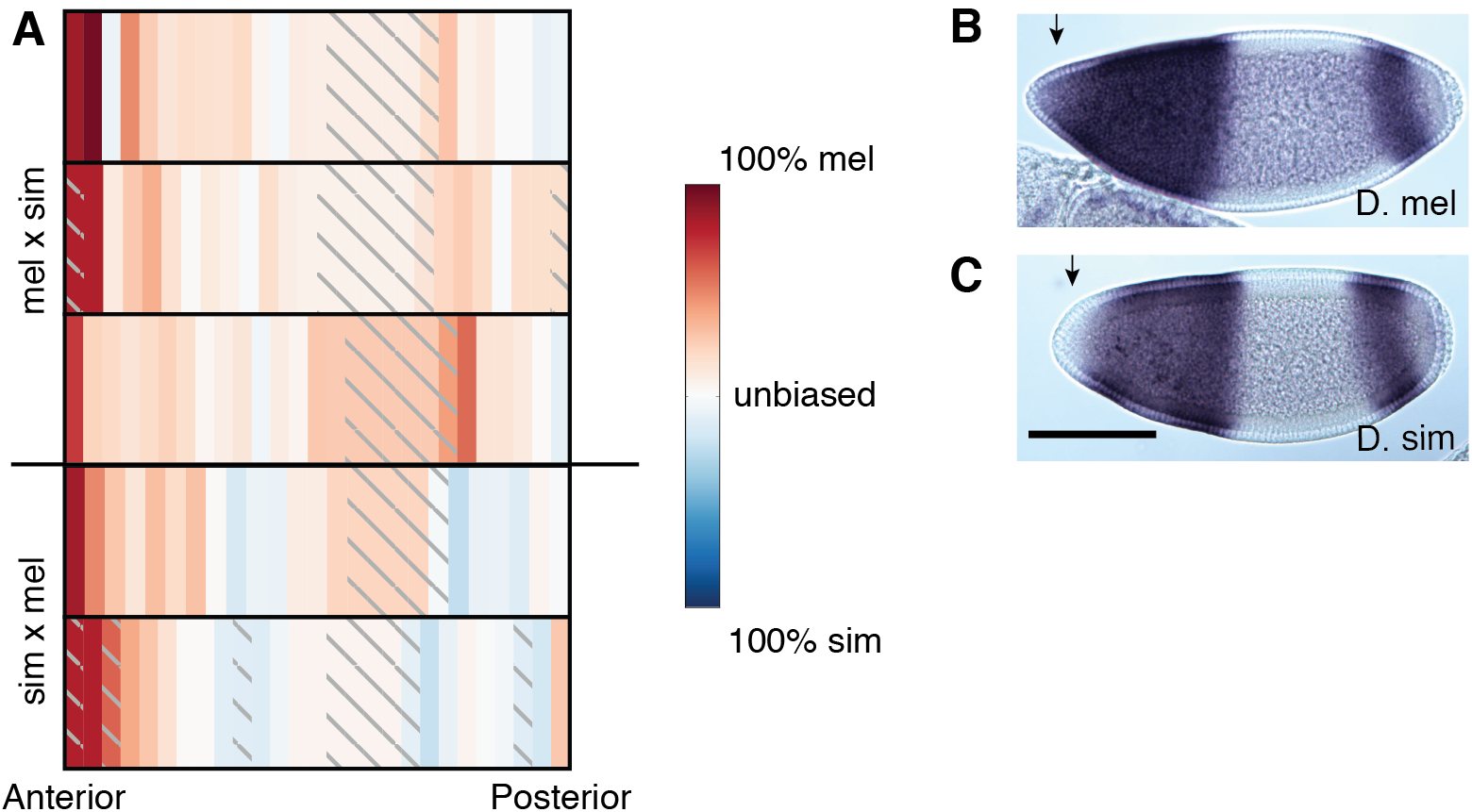
Hybrid embryos show strong melanogaster-specific expression of *hunchback* in the anterior. A) Heatmap of svASE of *hb* shows a significant *D. melanogaster* bias in the anterior tip of the embryo. Each row is a different embryo. Embryos with a melanogaster mother are above the horizontal line. B-C) *In situ* hybridization for *hb* in parental embryos. Images are arranged anterior to the left and dorsal up.

We next examined known regulatory sequences near *hb* for changes in TF binding sites that might cause the strong ASE in the anterior tip of the embryo. We downloaded from RedFly all known CRMs and reporter constructs with *hb* as a target (***Gallo et al., 2011***). There are three known minimal CRMs for *hb* that have been tested for embryonic activity using transgenic constructs: the canonical anterior CRM proximal to the *hb* promoter (***Driever and Nüsslein-Volhard, 1989***; ***Schröder et al., 1988***), a more distal “shadow” CRM (***Perry et al., 2011***), and an upstream CRM that drives expression in both the anterior and posterior domains, but not the anterior tip of *D. melanogaster* (***Margolis et al., 1995***) (***Figure 3A***). We excluded the upstream CRM from further consideration and used FIMO to scan the other regulatory sequences for motifs of the 14 TFs with ChIP signal near *hb* (***Li et al., 2008***; ***Bailey et al., 2015***). Binding in the canonical Bicoid-dependent anterior element gained only a single weak Bicoid motif in *D. simulans* relative to *D. melanogaster* (***Figure 3B***), and the distal “shadow” CRM gained Twist and Dichaete binding motifs between *D. melanogaster* and *D. simulans* (***Driever et al., 1989***; ***Perry et al., 2011***) (***Figure 3C***). Unsurprisingly, binding sites for other TFs outside the core regulatory elements displayed pervasive apparent turnover, with multiple gains and losses between the species (***Figure 3–Figure supplement 1***) (***Lusk and Eisen, 2010***; ***He et al., 2011***).

**Figure 3.**
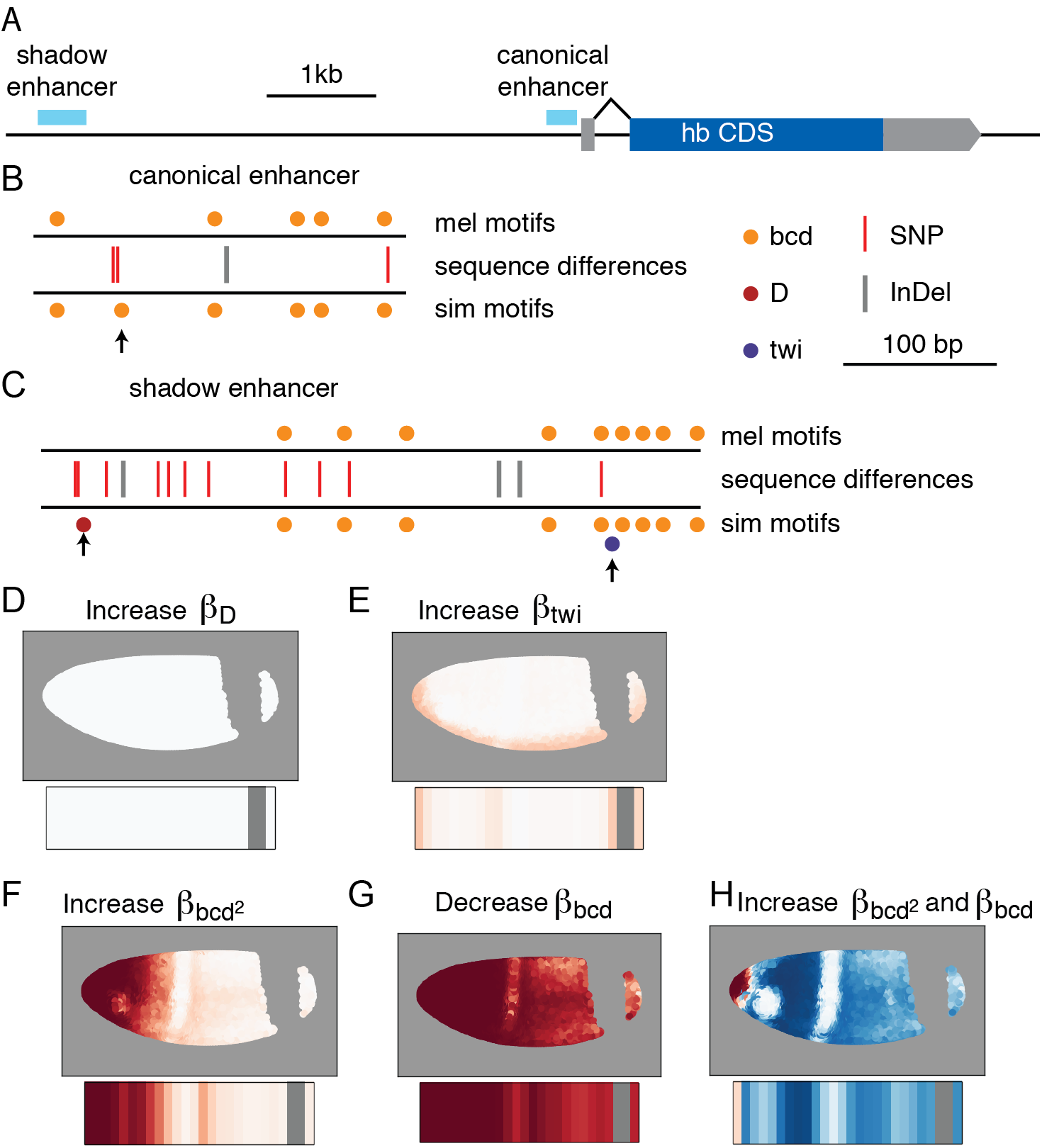
Cis-regulatory changes in *hb* regulatory regions could cause the observed svASE. A) Regulatory elements near the zygotic *hunchback* transcript. B-C) FIMO binding motifs and inter-specific variants of the anterior activator (B) and shadow CRM from ***Perry et al.*** (***2011***) (C). Species-specific predicted binding sites are highlighted with arrows. D-H) Predicted ASE from adjusting strength of each TF in the model in order to maximize the variance in the real ASE explained by the predicted ASE. Predicted ASE per nucleus is shown above and predicted ASE in a sliced embryo is shown below.

Anterior zygotic expression of *hb* is driven primarily by Bicoid, but there are details of the expression pattern at mid-stage 5 that cannot be explained by the relatively simple Bicoid gradient, and the loss of expression at the anterior tip of *D. simulans* cannot be explained by additional Bicoid activation. In order to more fully understand how this pattern might be specified and what the effects of binding site changes could be, we took a modeling-based approach similar to ***Ilsley et al.*** (***2013***). We used the 3-dimensional gene expression atlas from ***Fowlkes et al.*** (***2008***) to test regulators in a logistic model for the anterior *hunchback* expression domain (see Methods). The model included a linear term for every gap gene TF bound in the anterior activator CRM (***Li et al., 2008***) and a quadratic term for Bicoid to account for recent observations that it may lose its ability to act as an activator at high concentrations (***Fu and Ma, 2005***; ***Ilsley et al., 2013***). The best fit model (***Figure 3—data 1***) had the strongest coeicients for the two Bicoid terms, consistent with previous studies examining *hb* output as a simple function of Bcd concentration (***Driever et al., 1989***; ***Driever and Nüsslein-Volhard, 1989***; ***Gregor et al., 2007***). All the other TFs that bind to the locus are understood to be either repressors or have unclear direction of effect; consistent with this, most of the coeicients for those TFs are negative (***Reinitz and Levine, 1990***; ***Ganguly et al., 2005***; ***Small et al., 1991***). The exceptions to this are D and Twi which act as weak activators in the model, consistent with observations in the literature of bifunctionality for these TFs (***Aleksic et al., 2013***; ***Sandmann et al., 2007***).

We built this model to determine whether any of the binding site changes between *D. melanogaster* and *D. simulans* could plausibly explain the ASE that we observe in *hb*. Therefore, we did not make any effort to determine the minimal set of TFs that would drive the *hb* pattern, nor did we include a term to model predicted autoregulation (***Treisman and Desplan, 1989***; ***Holloway et al., 2011***).

In order to predict what effect the binding changes would have on expression in a *D. simulans* (or hybrid) embryo, we adjusted the coeicient for each TF independently to find the coeicient that best predicted the observed ASE. We then compared the output of the *D. melanogaster* model to the adjusted one (***Figure 3D***-H). Adjusting the Bcd coeicients, either alone or in tandem, produced the predicted ASE pattern most similar to the actual expression differences we observed between the species. We therefore hypothesized that the additional Bicoid site produced the smaller *D. simulans hb* anterior domain.

To test this prediction, we used CRISPR-Cas9 and homology-directed repair genome editing to introduce the Bicoid binding site SNPs from *D. simulans* into *D. melanogaster* embryos (***Gratz et al., 2014***; ***Port et al., 2014***). In order to avoid any transgene-specific ectopic staining, we edited the endogenous *hunchback* regulatory locus in *D. melanogaster*. We created 2 homozygous lines based on separate integration events, but with identical *D. simulans* sequence at the *hb* regulatory locus. We then tested these lines using *in situ* hybridization, and found that edited lines lose *hb* expression in the anterior tip, making the pattern much more similar to *D. simulans* (***Figure 4*** and ***Figure 4***—Figure supplement 1).

**Figure 4.**
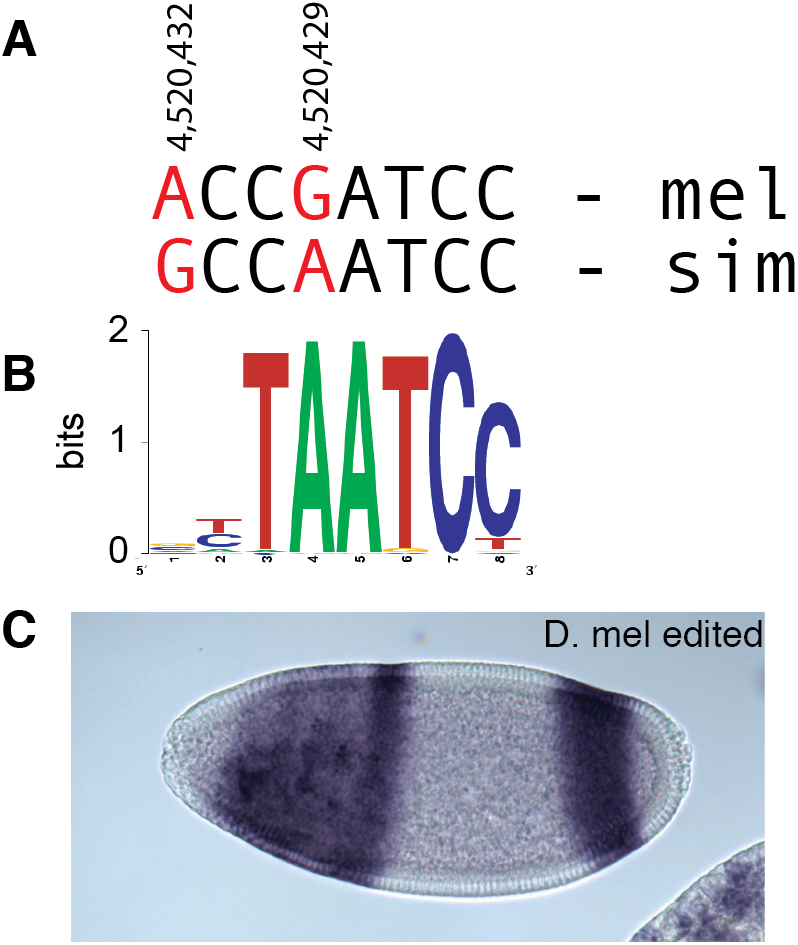
CRISPR-Cas9-mediated editing shows a Bicoid site in *D. simulans* is responsible for the change in expression pattern. A) A pair of SNPs in the canonical *hb* CRM at the indicated coordinates on *D. melanogaster* chromosome 3R. SNPs between *D. melanogaster* and *D. simulans* marked in red. B) The Bicoid binding motif. C) Representative *in situ* hybridization image for *hb* in a *D. melanogaster* embryo with the two base-pairs altered to match the *D. simulans* sequence at the canonical CRM.

We noticed that of the two SNPs that differ between *D. melanogaster* and *D. simulans*, the SNP that is outside of the core Bicoid binding motif is fixed in a survey of 20 *D. simulans* lines, whereas the SNP within the core of the motif (position 4,520,429; ***Figure 4A***) is segregating in *D. simulans* and is the minor allele (present in only 3 of the 20 lines in ***Rogers et al.*** (***2015***)). To test the role of this variant in isolation, we screened a number of *D. simulans* stocks and found a line, “sim 188” (***Machado et al., 2016***), that had the *D. melanogaster*-like sequence in the core of the Bicoid motif. When we performed *in situ* hybridization, we found that *hunchback* expression was present at the anterior tip of the embryo (***Figure 4–Figure supplement 2***), as in *D. melanogaster*, lending further strength to the hypothesis that the difference in expression pattern is due to Bicoid binding, and that the core Bicoid motif SNP is primarily responsible.

## Discussion

The study of allele-specific expression in F1 hybrids is a powerful tool for probing the evolution of gene expression (***Fraser, 2011***; ***Wittkopp and Kalay, 2012***). However, previous studies on *Drosophila* hybrids have been limited in their ability to pinpoint the causal variants responsible for the observed cis-regulatory divergence (***Wittkopp et al., 2004***; ***Graze et al., 2009***; ***Coolon et al., 2014***). In particular, the use of adult samples comprising multiple cell types meant that there was comparatively little information about the regulatory environment. In contrast, by focusing on the *Drosophila* embryo and using spatially-resolved samples, we were able to leverage decades of genetic and functional genomic information in *D. melanogaster* (***Driever and Nüsslein-Volhard, 1989***; ***Tomancak et al., 2007***; ***Li et al., 2008***; ***Fowlkes et al., 2008***; ***Gallo et al., 2011***; ***Li et al., 2011***; ***Shazman et al., 2014***). Combining this information with mathematical modeling of gene expression patterns yielded specific, testable predictions about which sequence changes produced the observed expression differences (***Figure 3***). Finally, by using CRISPR-mediated genome editing, we were able to directly confirm the genetic basis of the divergence in *hb* expression.

Although we were careful to minimize mapping bias in the detection of ASE, it is possible that non-zero ASE in any given gene is due to purely technical effects. However, by comparing parts of the same embryo to one another, we can effectively control for technical effects; even if the absolute level of ASE is incorrect, the variation is still meaningful. More importantly, changes in the position but not the absolute level of expression would be lost in bulk samples, and spatially restricted expression changes would tend to be washed out by more highly expressed and less variable regions.

A previous study found allele-specific expression for Ì 15% of genes in a *D. melanogaster* ◊ *D. simulans* hybrid adult ***Coolon et al.*** (***2014***). Considering that 400-600 genes have AP expression patterns in blastoderm stage embryos (***Tomancak et al., 2007***; ***Combs and Eisen, 2013***), our results suggest a roughly similar fraction of these patterned genes have strong svASE. We chose to restrict our study to the AP axis because it is straightforward to generate well-aligned slices with the long axis of a prolate object; there are no doubt many genes with dorsal-ventral expression differences as well, especially since DV CRMs tend to be shorter (***Li and Wunderlich, 2017***), and thus potentially more sensitive to sequence perturbation than AP CRMs.

Our experiment with editing the *hunchback* locus also suggested that Bicoid loses its activator activity at the anterior tip of the embryo. Although ***Ilsley et al.*** (***2013***) found that the two Bicoid terms have a net negative effect in the anterior tip of the embryo for *eve*, in our model the balance of the linear activation term and the quadratic repression term is such that at the anterior tip the two approximately cancel each other out. This is consistent with the observations that Torso signaling phosphorylates Bicoid in the anterior and deactivates it (***Ronchi et al., 1993***; ***Janody et al., 2000***), rather than making Bicoid function as a transcriptional repressor. On the other hand, despite lacking evidence that Bicoid can act as a repressor, the unmodified shadow CRM (which can drive expression in the anterior tip) is evidently not able to compensate for the reduced activity in the primary *D. simulans* CRM. Nor is it obvious that increased binding of an inactive factor would reduce expression.

We were not able to detect any aberrant phenotype of the altered *D. melanogaster* embryos engineered to have the *D. simulans hunchback* expression pattern. This is not surprising—although there are a number of subtle morphological, behavior, and physiological differences between *D. melanogaster* and *D. simulans* (***Orgogozo and Stern, 2009***), they are nevertheless generally regarded as indistinguishable as adults (***McNamee and Dytham, 1993***). Development is robust to large variation in the amount of *hunchback*, with hemizygous embryos giving rise to phenotypically normal adults (***Yu and Small, 2008***). Similarly, although embryos with varying Bicoid concentrations have widespread transcriptional changes, development is able to buffer these changes, at least in part due to differential apoptosis at later stages (***Driever and Nüsslein-Volhard, 1988***; ***Liu et al., 2013***; ***Combs and Eisen, 2017***; ***Namba et al., 1997***). It is also possible that the reduced *hb* expression in *D. simulans* matters only in particular stress conditions, but given the similar cosmopolitan geographic distributions of *D. melanogaster* and *D. simulans*, it is not obvious what conditions those might be. We believe that the informed modeling approach we have taken can serve as a model for dissecting other cis-regulatory modules. Eight genes with clear svASE are present in the BDTNP expression atlas (***Fowlkes et al., 2008***), and preliminary modeling of the four genes without pair-rule-like striping patterns suggested plausible binding site changes that could be responsible (***Figure 3***—Figure supplement 4). In some of these cases, there are multiple binding site changes that could explain our observed svASE equally well, but predict different dorso-ventral gene expression patterns in *D. simulans*—in these cases, *in situ* hybridization for the gene with svASE should provide clearer hypotheses of the causal variants. This approach, when applied more broadly and in concert with evolutionary studies, should help refine our understanding of both the molecular mechanisms and phenotypic consequences of the evolution of spatial patterning.

## Materials and Methods

### Strains and hybrid generation

Unless otherwise indicated, we used DGRP-340 as the *D. melanogaster* strain, and w501 as the *D. simulans* strain. Males of both species were co-housed for 5 days at 18C in order to improve mating eiciency, then approximately twenty males were mated with ten 0–1 day old virgin females of the opposite species per vial with the stopper pressed almost to the bottom. After cohousing, males were sorted using eye color as a primary marker. 5 days later, flies from the vials with larvae were put into a miniature embryo collection cage with grape juice-agar plates and yeast paste (Genessee Scientific).

### RNA extraction, library preparation, and sequencing

We selected single embryos at the target stage (based on depth of membrane invagination) on a Zeiss Axioskop with a QImaging Retiga 6000 camera and transferred them to ethanol-cleaned Peel-a-way cryoslicing molds (Thermo Fisher). We then applied approximately 0.5 µL of methanol saturated with bromophenol blue (Fisher Biotech, Fair Lawn N.J.), then washed with clean methanol to remove the excess dye. Next, we covered the embryo in Tissue-Plus O.C.T Compound (Fisher Healthcare) and froze the embryo at -80 until slicing. We sliced the embryos using a Microm HM550 cryostat, with a fresh blade for each embryo to minimize contamination.

We used 1mL of TRIzol (Ambion) with 400 µg/mL of Glycogen (VWR) to extract RNA, ensuring that the flake of freezing medium was completely dissolved in the TRIzol. In order to determine the sex of each embryo, we generated cDNA from the RNA using SuperScript II (Invitrogen) and a gene-specific primer for Roc1a, which is on the X chromosome and has a 49bp *D. simulans* specific insertion. We then amplified bands (Primers: cca gat gga ggg agc agc ac(forward) and atc gcc cca cta gct taa gat ct (reverse) amplicon lengths: 99bp and 138 bp) to determine the sex of hybrid embryos.

Next, we randomized the order of the RNA samples (see Supplementary file 1), then prepared libraries using a slightly modified version of the SMART-seq2 protocol (***Picelli et al., 2014***). As described in ***Combs and Eisen*** (***2015***), instead of steps 2-5 of the protocol in ***Picelli et al.*** (***2014***), we added 1µL of oligo-dT and 3.7µL of dNTP mix per 10µL of purified RNA; in step 14, we reduced the pre-amplification to 10 cycles; from step 28 onwards, we reduced the volume of all reagents by five-fold; and at step 33, we used 11 PCR amplification cycles.

We sequenced libraries in 4 separate lanes on either an Illumina HiSeq 4000 or an Illumina NextSeq (See Supplementary file 1 for lane and index details).

### Sequencing data processing and ASE calling

In order to call mappable SNPs between the species, we used Bowtie 2 (***Langmead and Salzberg, 2012***, version 2.2.5, arguments very-sensitive) to map previously published genomic sequencing data for the lines in this study (SRR835939, SRR520334 from ***Mackay et al., 2012***; ***Hu et al., 2013***) onto the FlyBase R5.57 genome. We then used GATK (***DePristo et al., 2011***, version 3.4-46, arguments <m>-T HaplotypeCaller -genotyping_mode DISCOVERY –output_mode EMIT_ALL_SITES-stand_emit_conf 10 -stand_call_conf 30) to call SNPs.

Next, we created a version of the *D. melanogaster* genome with all SNPs that are different between the two species masked. We used STAR (***Dobin et al., 2013***, version 2.4.2a, arguments `–clip5pNbases 6`) to map each sliced RNA-seq sample to the masked genome. We further filtered our list of SNPs to those for which, across all the RNA-seq samples, there were at least 10 reads that supported each allele. We also implemented a filtering step for reads that did not remap to the same location upon computationally reassigning each SNP in a read to the other parent as described in ***van de Geijn et al.*** (***2015***).

To call ASE for each sample, we used the GetGeneASEbyReads script in the ASEr package (Manuscript in preparation, available at https://github.com/TheFraserLab/ASEr/, commit cfe619c69). Briefly, each read is assigned to the genome whose SNP alleles it matches. Reads are discarded as ambiguous if there are no SNPs, if there are alleles from both parents, or if the allele at a SNP does not match either parent. Additionally, for most subsequent analyses, ASE is ignored if the gene is on the X chromosome and the slice came from a male embryo (which only have an X chromosome from their mother). All other analysis scripts are available at https://github.com/TheFraserLab/HybridSliceSeq (commit c244b87).

### Earth Mover Distance and Spatial Patterning Diferences

Earth mover distance (EMD), as described in ***Rubner et al.*** (***1998***), is a non-parametric metric that compares two distributions of data in a way that roughly captures intuitive notions of similarity. It represents the minimal amount of work (defined as the amount moved multiplied by the distance carried) that must be done to make one pattern equivalent to another, as if transporting dirt from one pile to another. For each slice, we calculate the absolute expression of each gene using cuffinks v.2.2.1 (***Trapnell et al., 2013***). We normalize all absolute expression patterns by first adding a constant amount to mitigate noise in lowly expressed genes, and then by dividing by the total amount of expression in an embryo.

To compare between the hybrids and the parental embryos, we first calculated a spline fit for each gene on each of the parental embryos separately, first smoothing by taking a rolling average of 3 slices. We then fit a univariate spline onto the smoothed data using the Scipy “interpolate” package. Then, we recalculated the predicted expression for a hypothetical 27-slice embryo of each parent, then averaged the expression data. We next calculated the EMD between this simulated averaged embryo and each of the hybrid embryos. For each gene, we then performed a one-sided t-test to determine whether the hybrid embryos were more different from the average than the EMD between the parental embryos. Although 342 genes had a nominal p-value < .05, none of these remained significant after Benjamini-Hochberg multiple hypothesis testing correction.

To compare embryos between directions of the cross, we calculated the pairwise EMD between embryos within a direction of a cross (i.e. the three possible pairs of hybrid embryos with a *D. melanogaster* mother and the pair of embryos with the *D. simulans* mother) and the pairwise EMD between hybrid embryos with different parents (e.g. the first replicate of embryos). We then used a one-sided t-test to determine whether the EMDs were larger between groups than within. Benjamini-Hochberg FDR estimation yielded 171 genes with a q-value less than .05, whereas Bonferroni p-value correction yielded 12 genes at α.05.

### Identification of allele-specific expression patterns

In order to call a gene as strongly biased, we required that gene have at least 10 slices with detectable ASE, with at least 65% of those slices having at least 66.7% of reads from the same parent (maternal, *D. simulans*, or *D. melanogaster*, as appropriate). To detect genes with more subtle, yet consistent, overall ASE we summed the ASE scores for each embryo separately. To create a null distribution, we randomly flipped the sign of each ASE score then summed the ASE of the randomized matrix, repeating 50,000 times. We then combined the p-values from each embryo using Fisher’s method, ignoring scores from X-chromosomal genes in male embryos. To estimate a false discovery rate, we compared the number of genes with a given p-value to the number expected at that p-value under a uniform distribution.

To call svASE, we fit a 4-variable least-squares regression of either a sigmoidal logistic function (*f* (*x*)= *A*/(1 + exp(*w*(*x* − *x*_0_))) − *y*_0_) or a peak-like Gaussian function (*f* (*x*)= *A*⋅exp(-(*x* - *x*_0_)^2^/*w*^2^) *y*_0_). We then consi<u>de</u>red any gene where the fit explained at least <u>4</u>5% of the variance 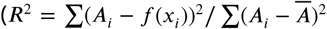, where *A*_*i*_ is the ASE value in the *i*th slice, and 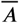 is the average ASE value for that gene) as having svASE.

To calculate a false discovery rate, we shuffed the columns (i.e. the spatial coordinates) of the ASE matrix 1,000 times. For each of the shuffes, we fit both of the ASE functions. Most of the shuffed matrices yielded no fits that explained at least 45% of the variance, only a handful of the matrices yielded a single gene that cleared the threshold, and no shuffed matrix had two or more genes that cleared the threshold.

### Spatially varying splicing diferences

To look for overall spatially varying splicing differences, we used the DEX-seq script `prepare_annotation` to identify exonic parts (***Anders et al., 2012***). For each exonic part in each slice, we calculated percent spliced in (PSI) (***Schafer et al., 2015***). Then we followed the same fitting procedure as for the allele-specific expression, with the same cutoff of 45% of the variance explained by the fit.

To look for spatially varying, allele-specific splicing, we adapted the ideas of ***Li et al.*** (***2016***) to look specifically at reads that support a splice junction. We used the LeafCutter script `leafcutter_cluster` on all of the mapped, de-duplicated reads to identify splice junctions that have at least 50 reads across our entire dataset. Then, for each read mapping to each well-supported splice junction, we used a custom script to assign it to either *D. melanogaster* or *D. simulans* as above. We then calculated an allele-specific splicing preference index as in equation 1 above. Finally, we used the same fitting procedure as above, except we used a relaxed cutoff of 25% of the variance explained, since only 1 gene had greater than 45% of its variance explained by a fit.

### Identification of binding site changes and predicted efects on hybrid embryos

For *hunchback* we used the coordinates for the regulatory elements as defined in the RedFly database to extract the sequence of each regulatory region from the reference sequence files (***Gallo et al., 2011***). For the other genes whose regulatory programs we investigated for causal binding changes, we used Bedtools to find any non-exonic DNase accessible region within 15,000 bp of each gene (***Quinlan and Hall, 2010***; ***Thomas et al., 2011***). We then used BLAST v2.3.0+ to search for the orthologous region in *D. simulans*. We combined motifs from the databases in ***Shazman et al.*** (***2014***); ***Enuameh et al.*** (***2013***); ***Kulakovskiy et al.*** (***2009***); ***Kulakovskiy and Makeev*** (***2009***); ***Bergman et al.*** (***2005***) by taking the most strongly-supported motif for a given TF, then we used the FIMO tool of the MEME suite to search for binding sites for all TFs with known spatial patterns (***Grant et al., 2011***; ***Bailey et al., 2015***).

In order to construct a model of transcription regulation for the other genes with svASE and simple expression patterns in the ***Fowlkes et al.*** (***2008***) atlas, we built models that contained the TFs with binding changes for the target gene as well as up to 4 other TFs with localization data in the ***Fowlkes et al.*** (***2008***) atlas and known roles as patterning factors during early development (i.e. Bcd, Gt, Kr, *cad, tll, D, da, dl, mad, med, shn, sna, twi, zen, brk, emc, numb, rho, tkv* and *Doc2*); when available, we used protein localizations instead of RNA *in situ* hybridization (i.e. for Bcd, Gt, and Kr). For a given combination of factors, we used the Python Statsmodels package to fita logistic regression to the anterior stripe of *hunchback* (***Seabold and Perktold, 2010***). In line with the procedure in ***Ilsley et al.*** (***2013***), we separated the two *hunchback* expression domains and fit the data on nuclei with either the anterior stripe or no *hunchback* expression. We then selected the best model based on fraction of variance in the original data explained by the fit.

To estimate the likely effect of each transcription factor change, we adjusted the relevant parameter(s) in the model by a range of values (see ***Figure 3***—Figure supplement 3). We then generated predicted svASE by predicting expression in each nucleus under the original model and the model with the relevant parameter(s) changed, grouping the nuclei by x-coordinate to simulate slicing, then combining the expression of each nucleus *i* in each slice *s* in an analogous manner to equation 1:

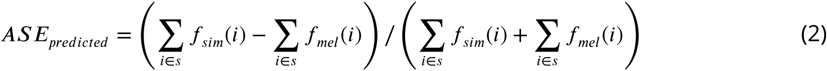

We then computed the Pearson correlation of the predicted and real ASE values and measured the fraction of the variance in the real ASE explained by the predicted ASE. In general, both measurement approaches suggested the same direction of change to the coeicient, although the absolute magnitude of change that yielded the “best” result may have been different.

### Genome Editing and Screening

We inserted the *D. simulans* SNPs into *D. melanogaster* using CRISPR-Cas9 directed cutting followed by homology directed repair (***Gratz et al., 2014***). We inserted the gRNA sequence GGT ACA GGT CGC GGA TCG GT into pU6-bbsI (a generous gift from Tim Mosca and Liqun Luo). We injected the plasmid and a 133bp ssDNA HDR template (IDT, San Diego, CA) into y[1] Mvas-Cas9ZH2A w[1118] embryos (Bloomington Stock #51323, BestGene Inc, Chino Hills, CA). The edited sequence affects a recognition sequence for the restriction enzymes BsiE1 and MspI (New England Biolabs) which specifically cut the *D. melanogaster* and *D. simulans* sequences, respectively. We screened putatively edited offspring by PCR amplifying a region around the *hunchback* anterior CRM (primers CGT CAA GGG ATT AGA TGG GC and CCC CAT AGA AAA CCG GTG GA) then cutting with each enzyme separately. Presumptively edited lines were then further screened via Sanger sequencing.

For the *in situ* hybridization, we generated DIG-labeled antisense RNA probes by first performing RT-PCR on *D. melanogaster hunchback* cDNA using primers with a T7 RNA polymerase handle (AAC ATC CAA AGG ACG AAA CG and TAA TAC GAC TCA CTA TAG GGA GA), then creating full-length probes with 2:1 DIG-labeled UTP to unlabeled UTP (***Weiszmann et al., 2009***). We then performed *in situ* hybridization in 2-4 hour old embryos of each strain according to a minimally modified, low-throughput version of the protocol in ***Weiszmann et al.*** (***2009***) (https://www.protocols.io/view/in-situ-hybridization-g7bbzin). Stained embryos were imaged on the Zeiss Axioskop above.

### Additional Files

RNA-seq data is available from the Gene Expression Omnibus with accession GSE102233.

**Supplementary data file 1** Summary description of library construction and sequencing information, including Nextera barcodes, sequencer type, and lane.

**Supplementary data file 2** Allele-specific expression matrix.

## Acknowledgements

We would like to thank Dmitri Petrov for the generous use of incubators and space for fly work. We thank the entire Fraser lab for discussions and advice.

---

**Figure 1–Figure supplement 1.**
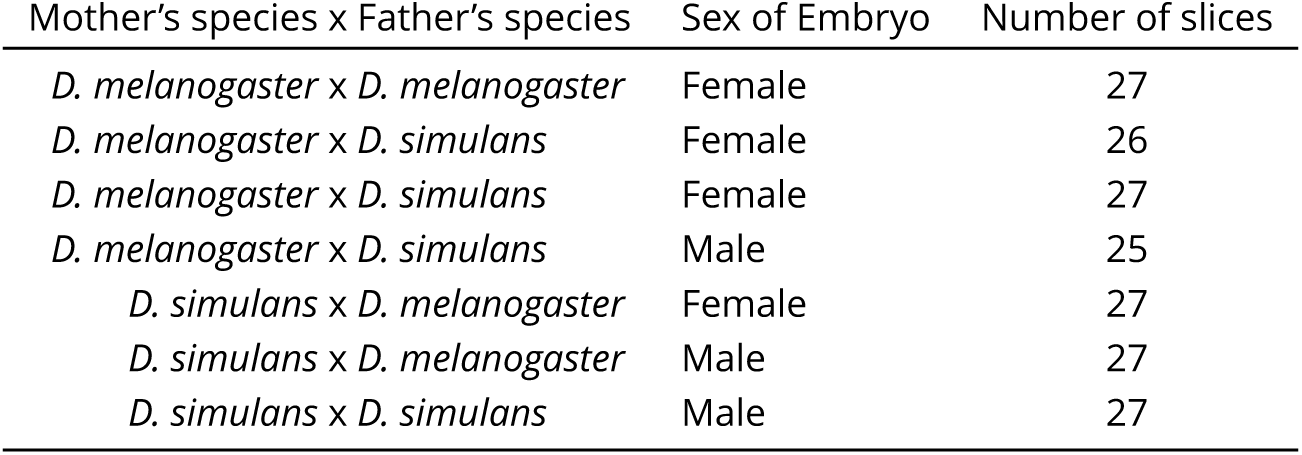
Summary data for embryos used.

**Figure 1–Figure supplement 2.**
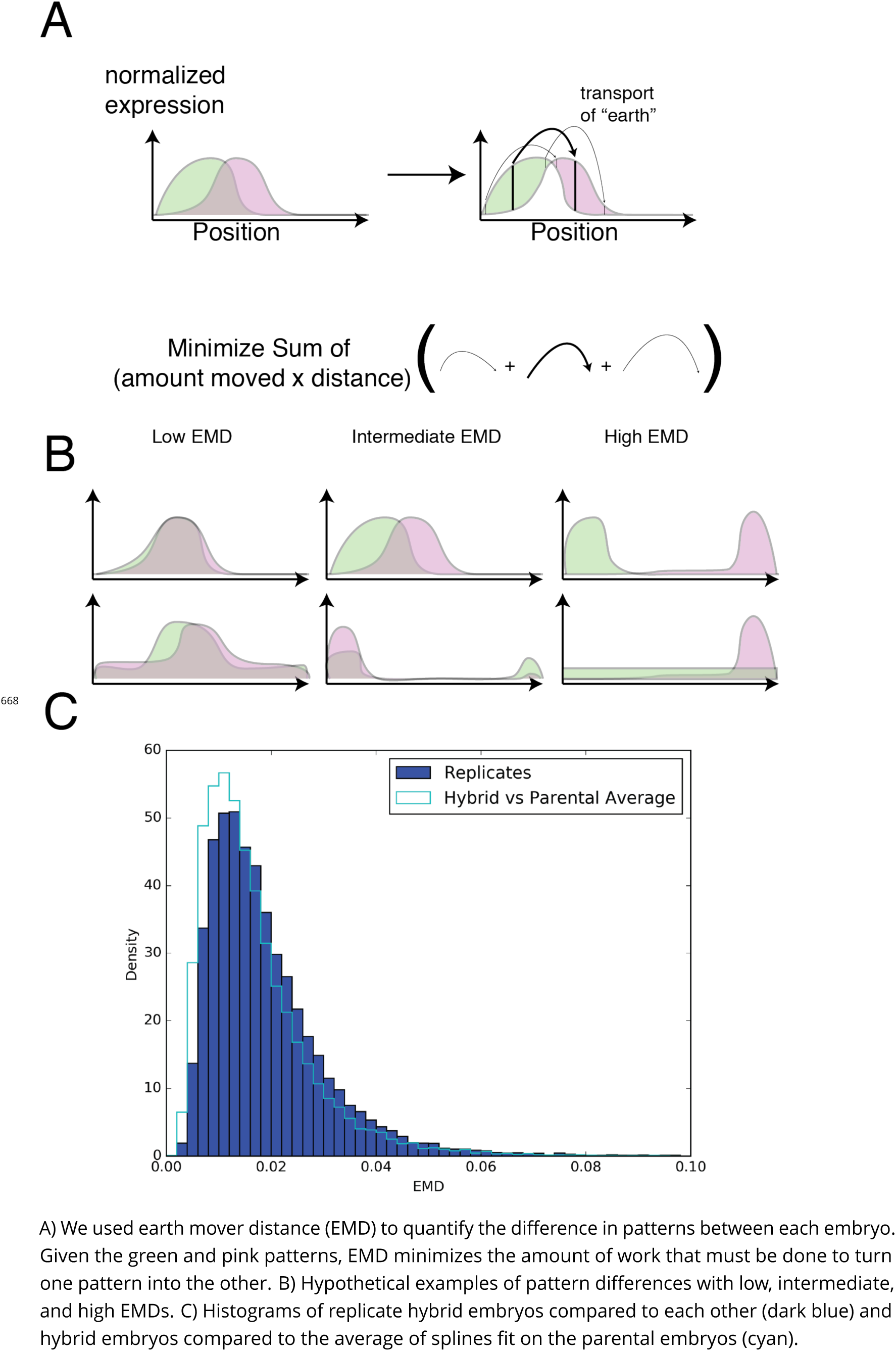
Using earth mover distance to identify genes with different expression patterns between the hybrids and the parents

**Figure 1–Figure supplement 3.**
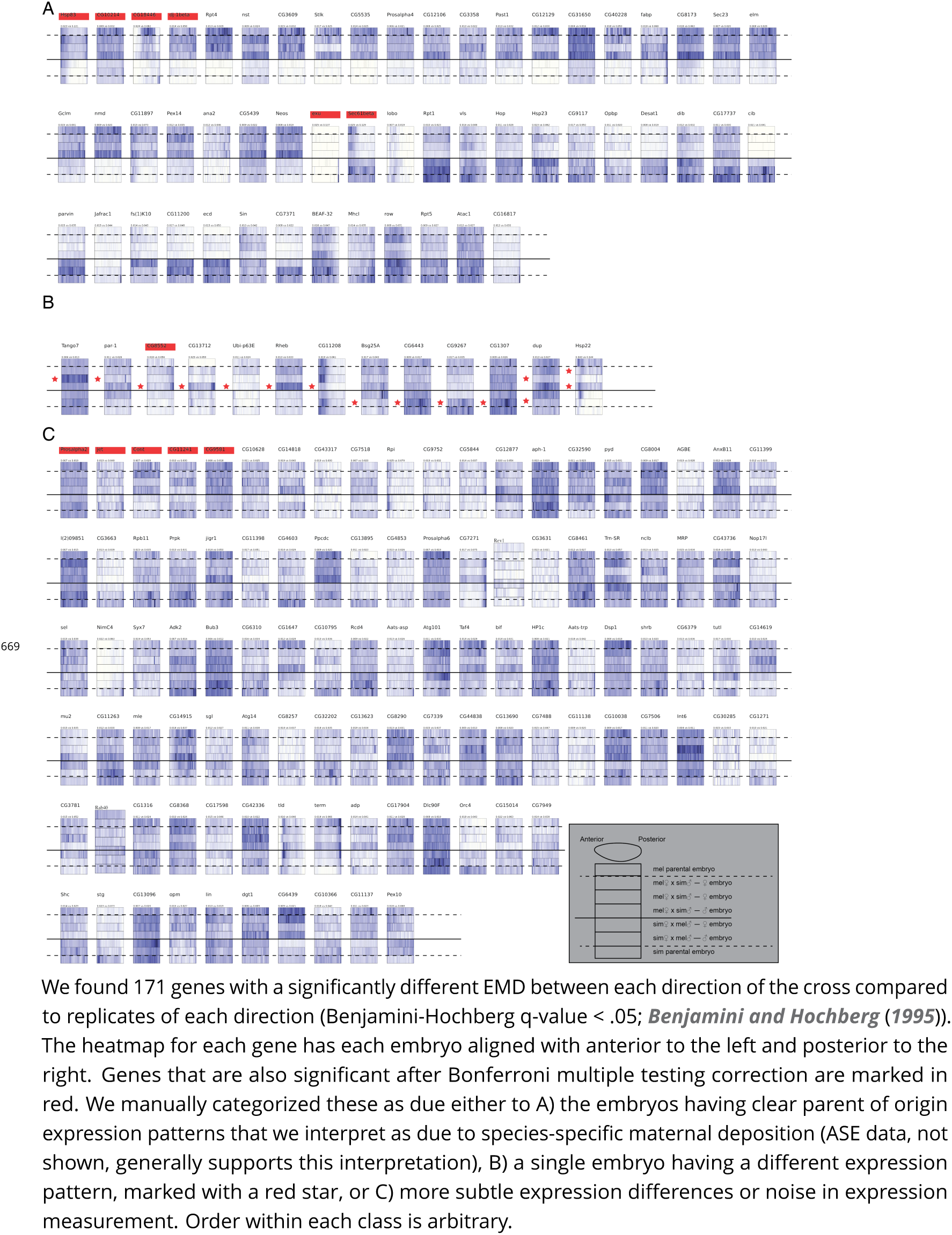
Using earth mover distance to identify genes with different expression patterns between the directions of the hybrid cross

**Figure 1–Figure supplement 4.**
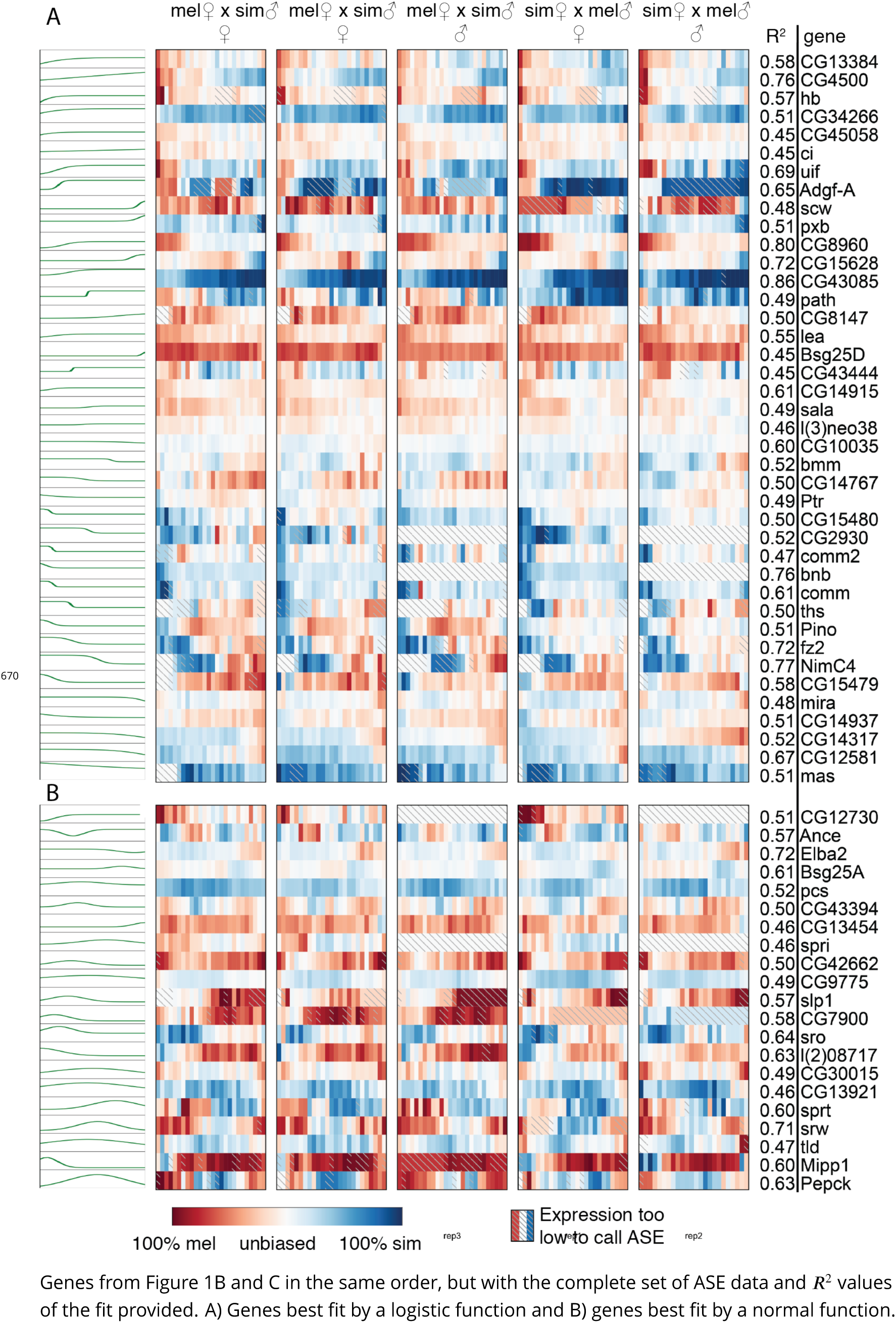
Complete heatmap of ASE for genes with svASE.

**Figure 1–Figure supplement 5.**
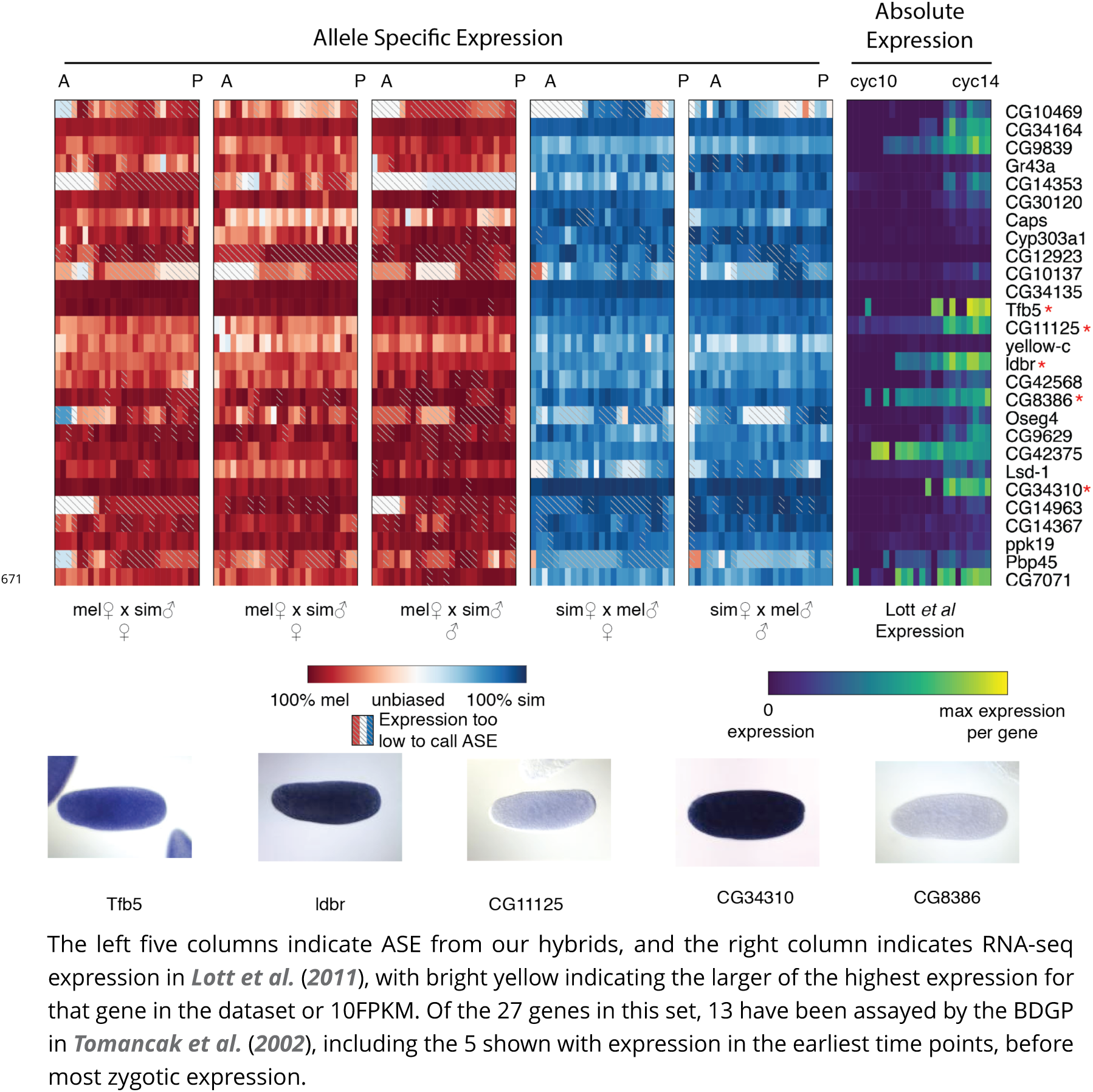
Genes identified as maternally deposited in our data but as zygotically expressed in ***Lott et al.*** (***2011***)

**Figure 1–Figure supplement 6.**
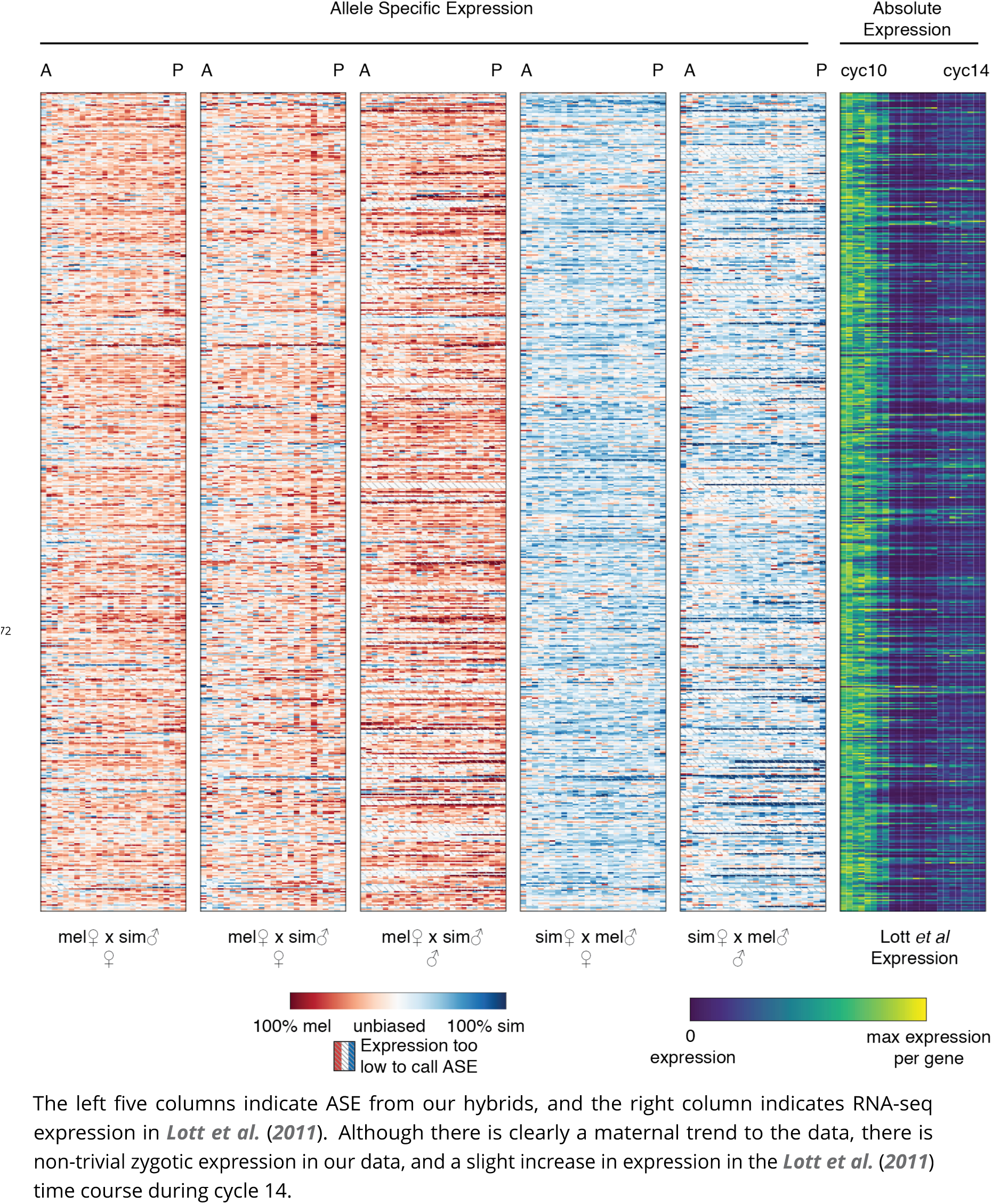
Genes identified as zygotically expressed in both crosses in our data but maternally deposited in ***Lott et al.*** (***2011***).

**Figure 1–Figure supplement 7.**
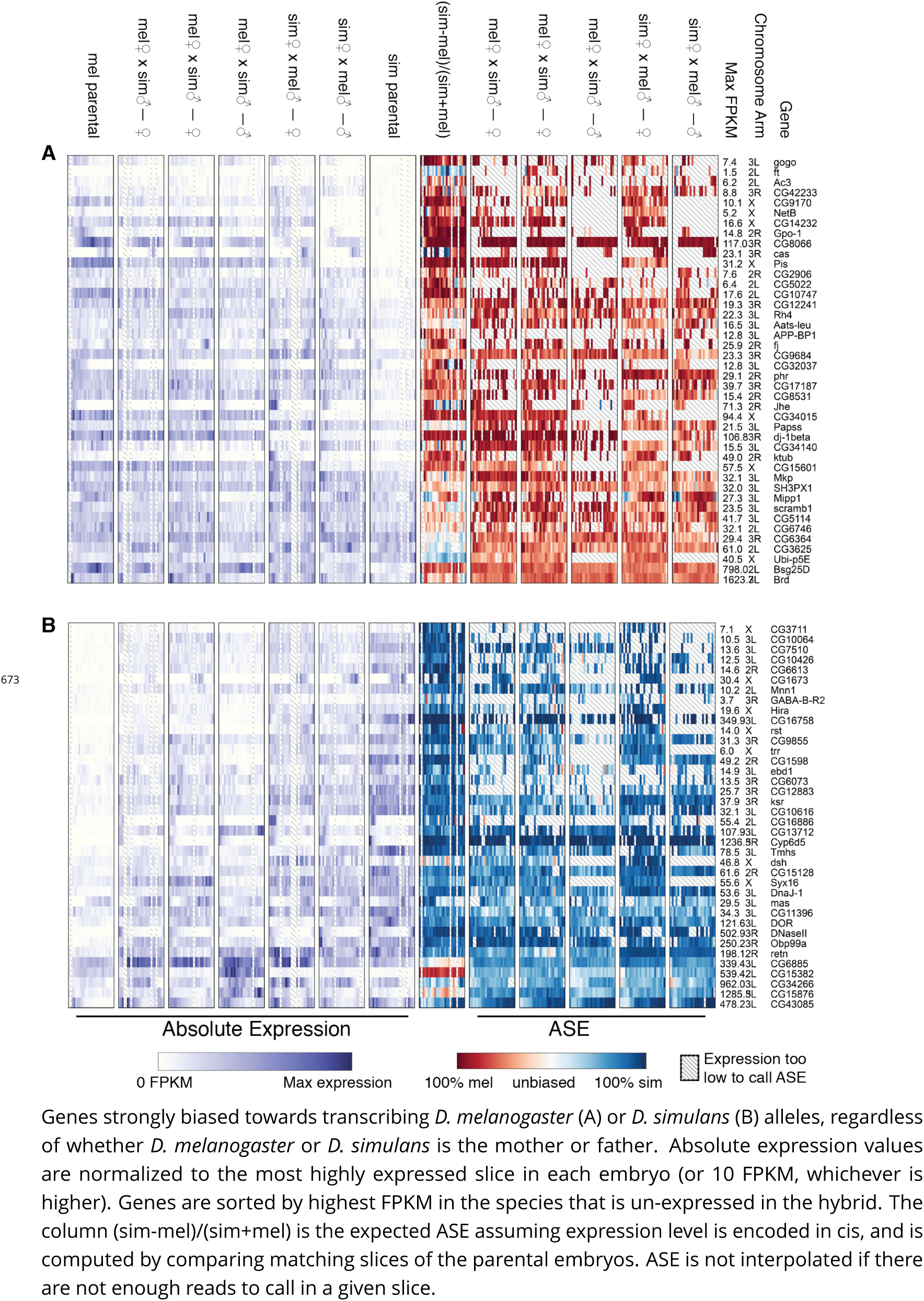
Genes with species-specific expression, regardless of parent of origin

**Figure 1–Figure supplement 8.**
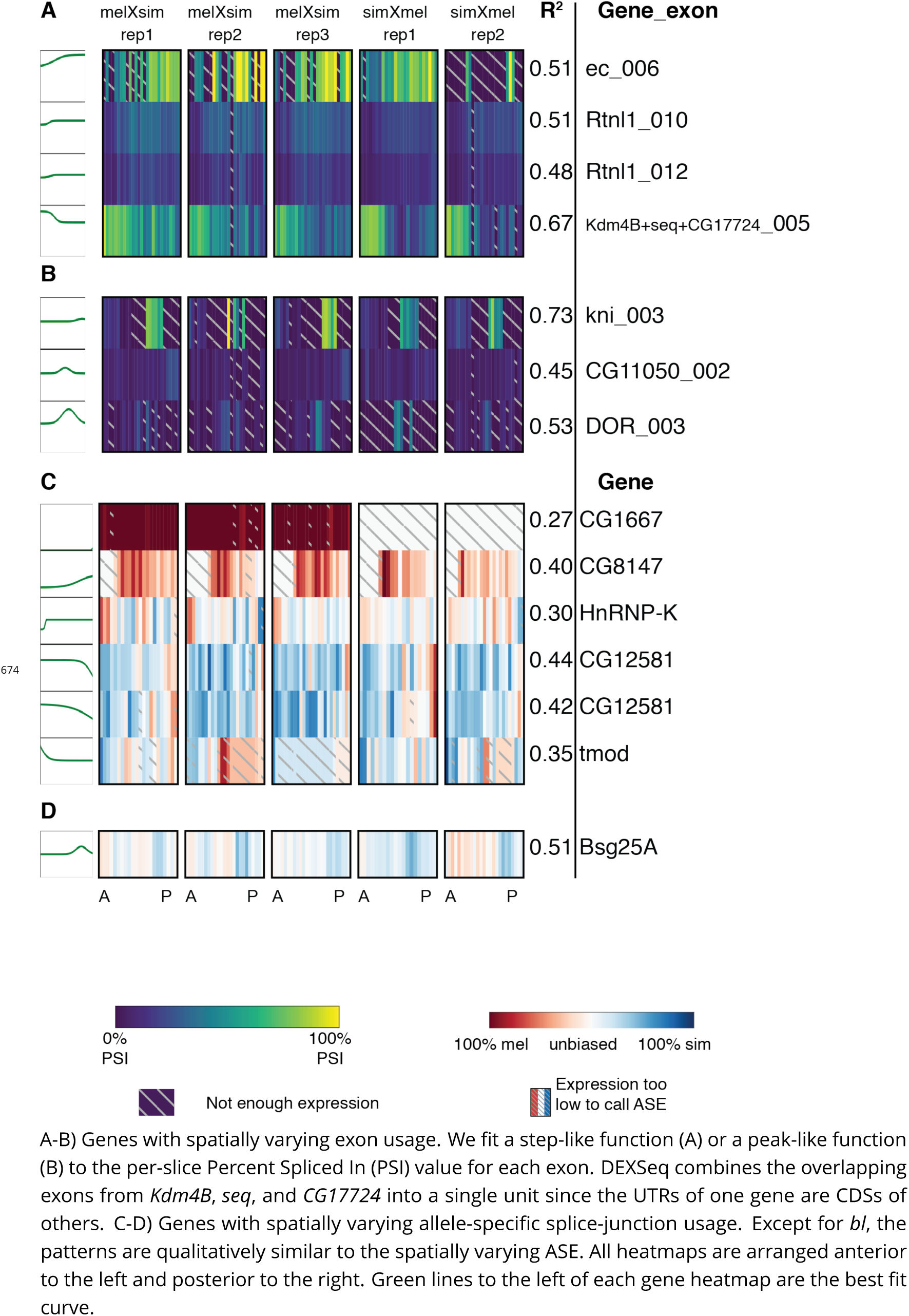
Genes with spatially varying splicing.

**Figure 3–Figure supplement 1.**
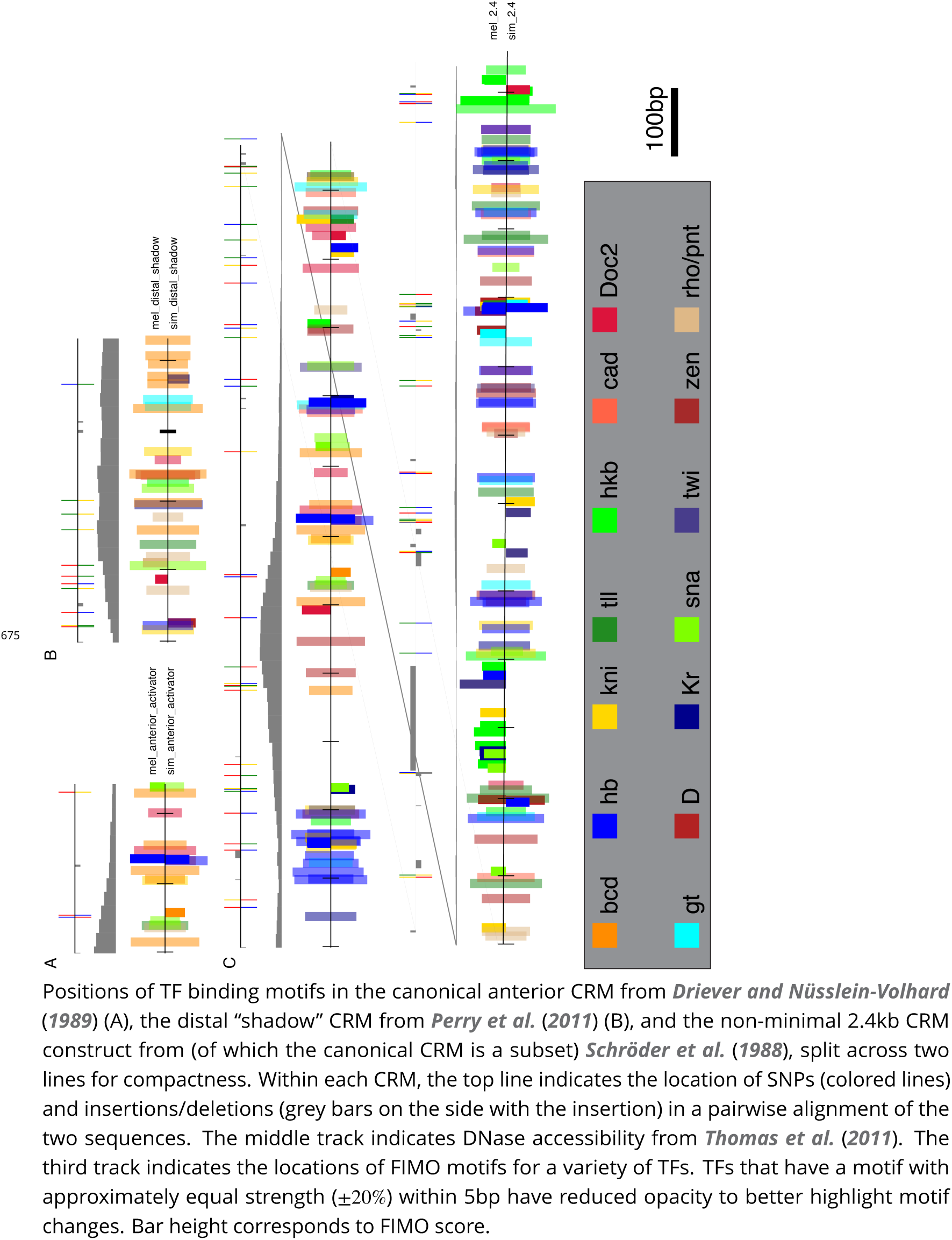
Motif content of the CRMs for all TFs included in the model.

**Figure 3–Figure supplement 2.**
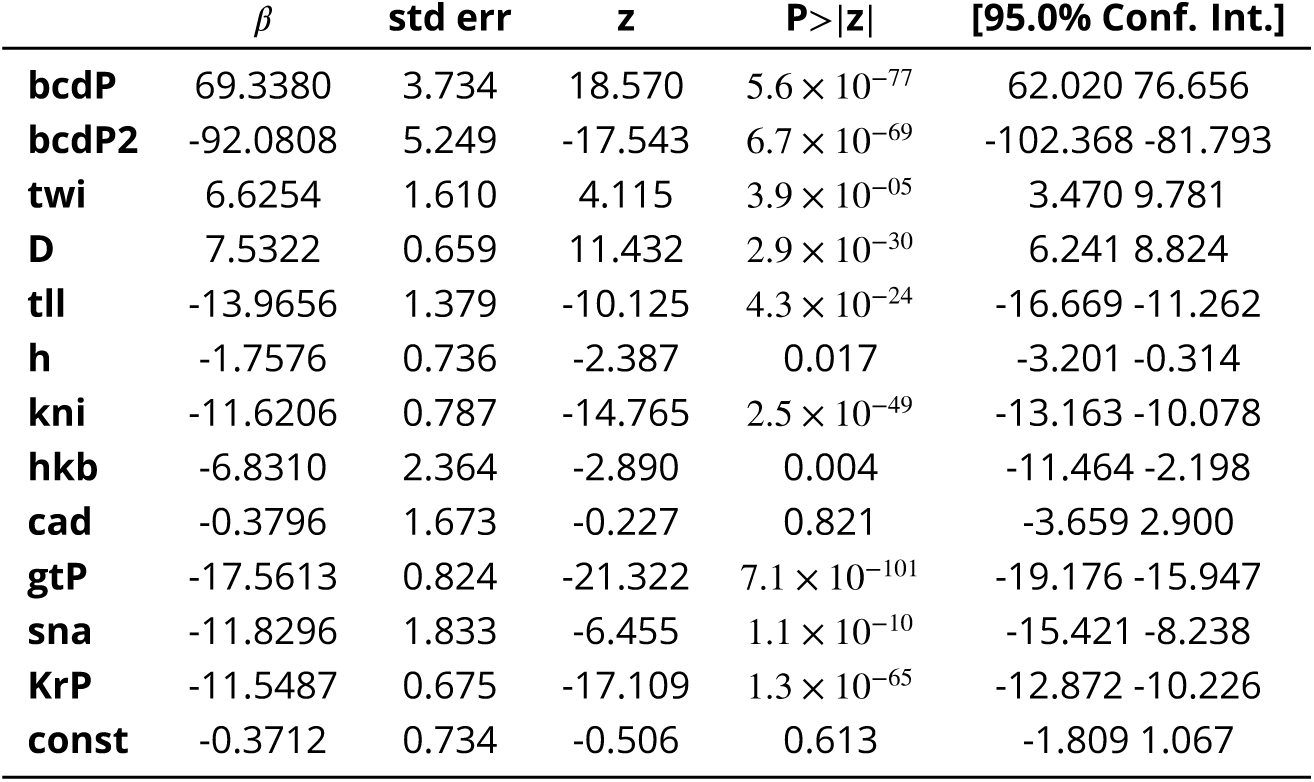
Coeicients of the best-fit model for TFs bound near the anterior activator of *hb*

**Figure 3–Figure supplement 3.**
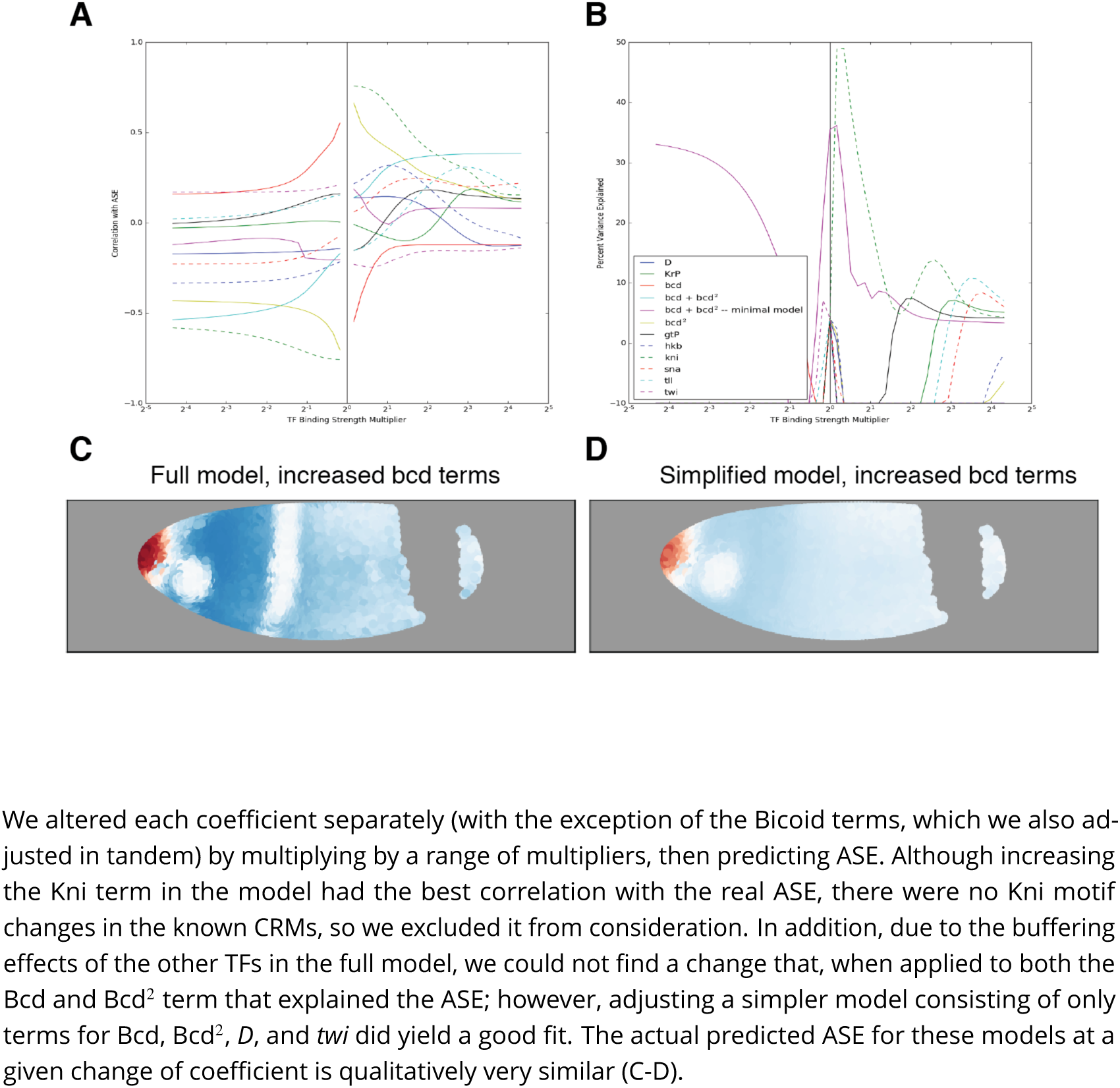
Correlation of the predicted *hb* ASE with the real ASE (A) and percent of the variance explained by predicted ASE (B) at a range of coeicient strengths.

**Figure 3–Figure supplement 4.**
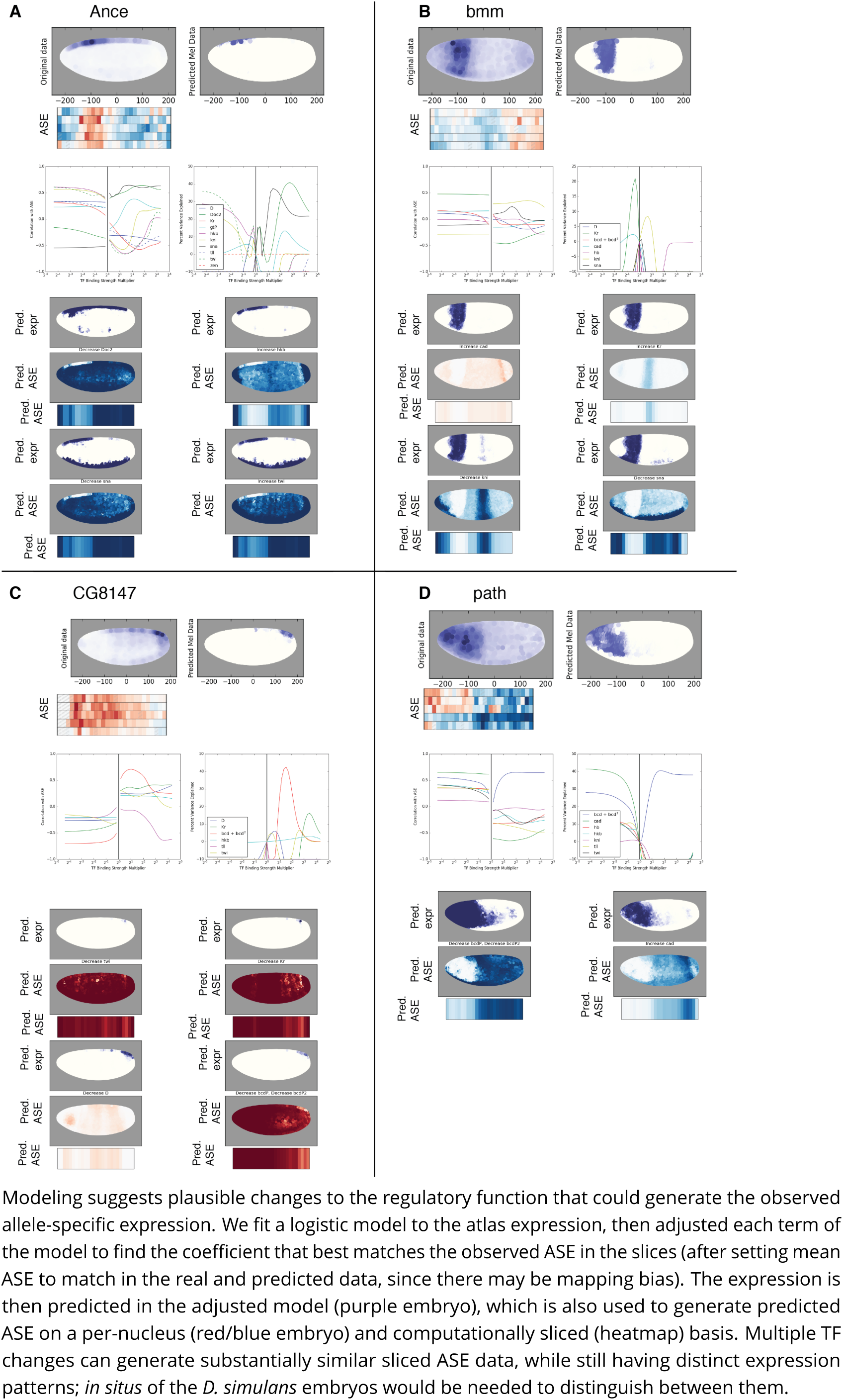
Proposed TF binding changes that generate svASE in *Ance*, *bmm*, *CG8147*, and *path*. We did not attempt modeling of the pair-rule genes *pxb*, *Bsg25A*, *comm2*, and *pxb*,

**Figure 4–Figure supplement 1.**
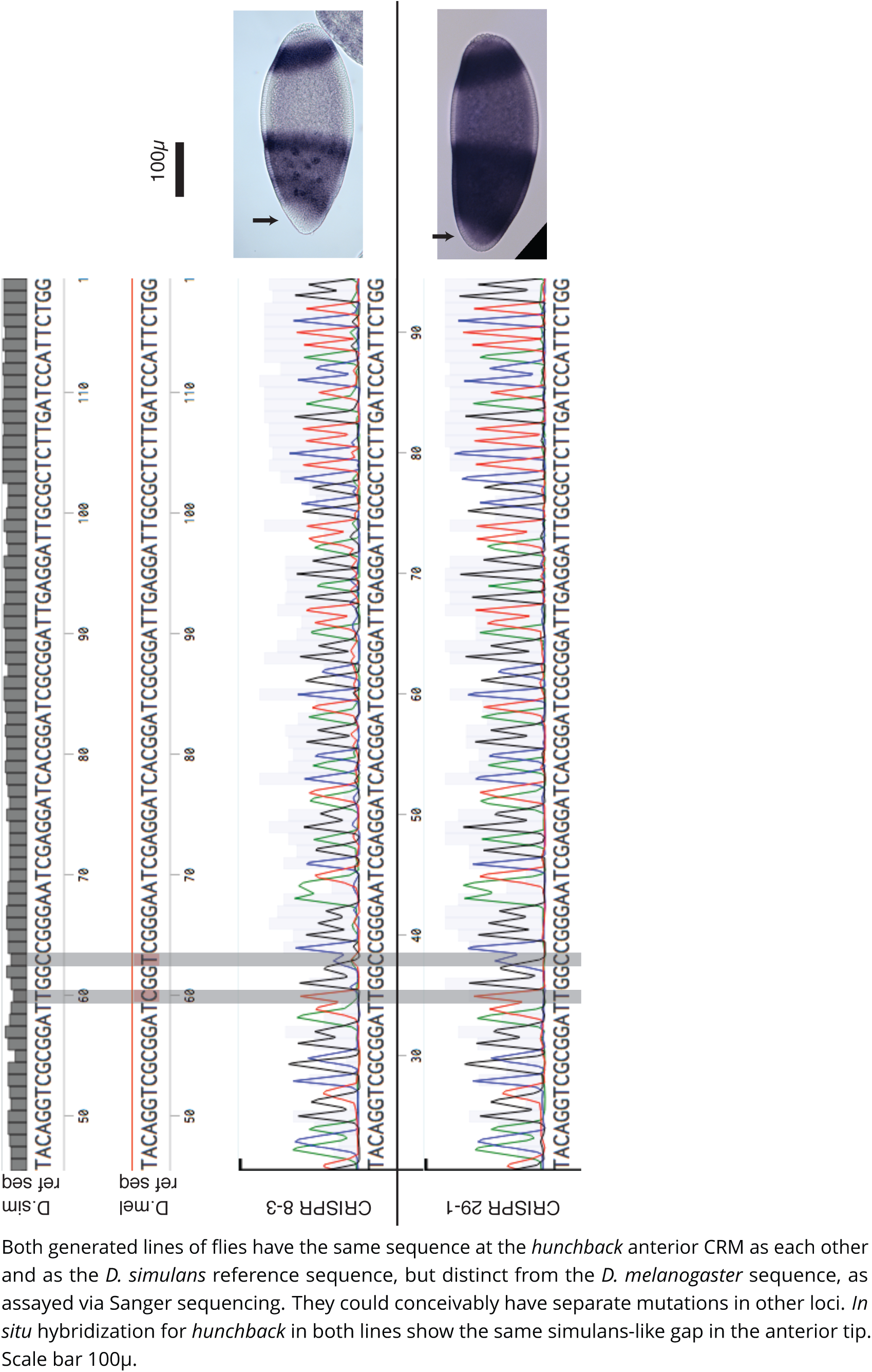
A second, independently edited *D. melanogaster* line also shows the anterior gap of hunchback expression

**Figure 4–Figure supplement 2.**
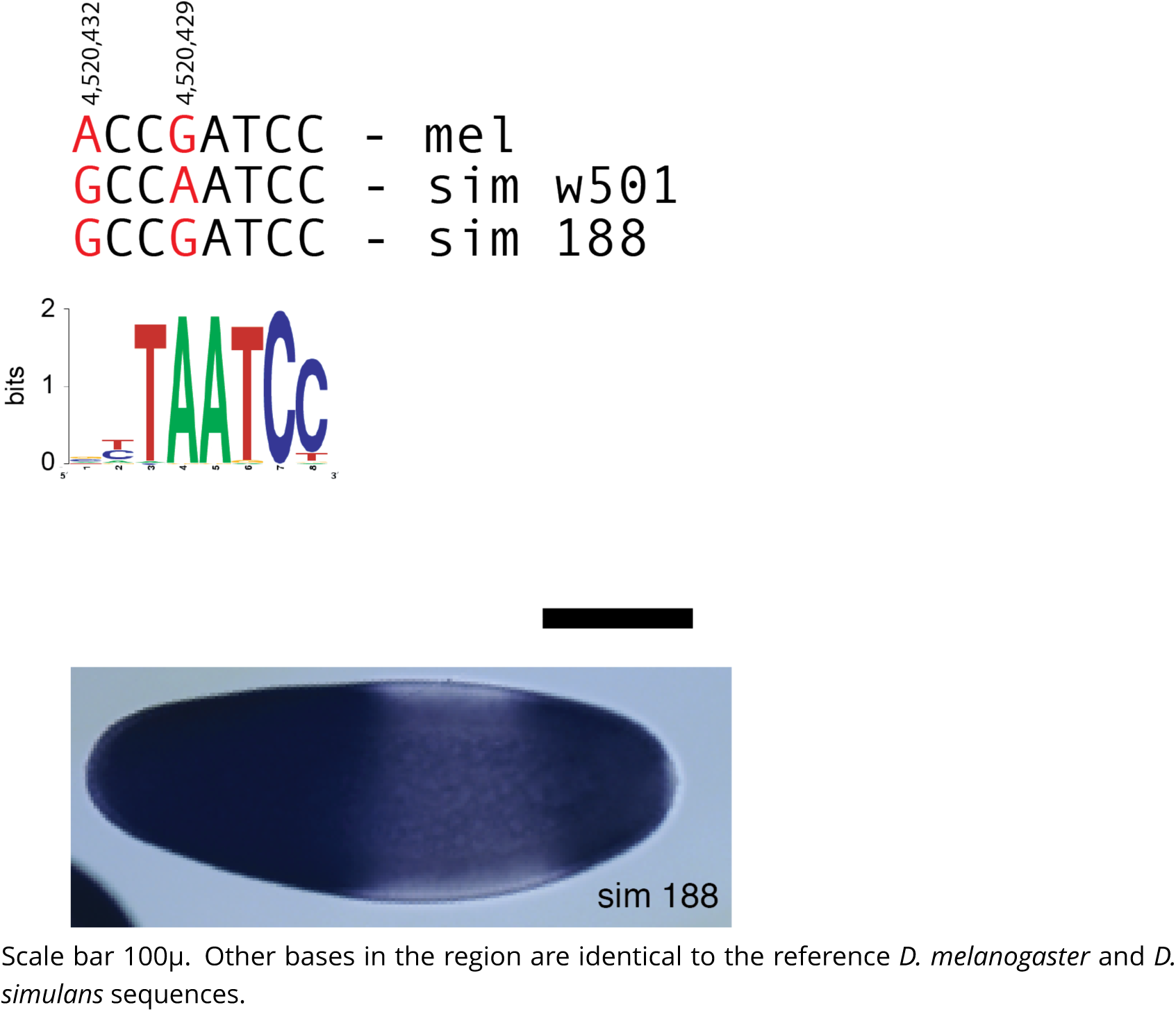
A naturally occurring strain of *D. simulans* with one of the base pair changes found in our edited line does not show the anterior gap of expression, closer to the *D. melanogaster* pattern.

